# Spatial pattern of serology against Borrelia and tick-borne encephalitis virus in roe deer *(Capreolus capreolus)*, Northern France, 2019-2022

**DOI:** 10.1101/2025.01.20.633885

**Authors:** Raphaëlle Métras, Matthieu Bastien, Valentin Ollivier, Armelle Poisson, Benoit Combes, Gaëlle Gonzalez, Teheipuaura Helle, Camille Migné, Ilona Eveline Suhanda, Younjung Kim, Hélène Verheyden, Thierry Boulinier

**Affiliations:** Sorbonne Université, INSERM, Institut Pierre Louis d’Épidémiologie et de Santé Publique (IPLESP), Paris, France; Entente de Lutte et d’Intervention contre les Zoonoses (ELIZ), Malzéville France; Entente Interdépartementale Rhône-Alpes de démoustication (EIRAD), Chindrieux, France; CEFE, UMR 5175, CNRS, University of Montpellier, EPHE, IRD, Montpellier, France; ANSES, INRAE, Ecole Nationale Vétérinaire d’Alfort, UMR VIROLOGIE, Laboratoire de Santé Animale, Maisons-Alfort, France; Université de Toulouse, INRAE, Comportement et Ecologie de la Faune Sauvage (CEFS), Castanet Tolosan, France

**Keywords:** Seroprevalence, Borrelia, tick-borne encephalitis, roe deer, spatial analysis

## Abstract

**Background:** Lyme borreliosis and tick-borne encephalitis (TBE) are tick-borne zoonoses of public health importance, transmitted in Western Europe by the same tick vector *Ixodes ricinus*. Roe deer (*Capreolus capreolus*) is a widespread feeding host for *I. ricinus* stages. Here, we explored the spatial extent of both pathogens in the wild in Northern France by assessing serological status in roe deer.

**Methods:** We collected 3,147 blood samples (2019-2022), and searched antibodies against both pathogens using ELISA tests. We fitted logistic models to investigate exposure to both or either pathogens, and explored spatial patterns using Kulldorf clustering scan tests.

**Results:** The overall ELISA IgG prevalence was 23.24% (95%CI [21.77-34.72]) for Borrelia and 8.64% (95%CI [7.45-9.86]) for TBEV. The median Borrelia seroprevalence varied from 4% to 42% between departments, with two high-risk clusters of exposure identified. TBEV seroprevalence did not show any spatial pattern, and positives by seroneutralisation tests were reported in eight departments.

**Discussion:** Our findings suggest the presence of both pathogens over large areas. Wider spatial sampling including areas at lower risk and improved serological methods would be beneficial to further validate those estimates.

## Introduction

Lyme borreliosis (LB) and tick-borne encephalitis (TBE) are tick-borne zoonoses of concern in temperate areas of the world. Caused by the infection with *Borrelia burgdorferi sensu lato* (Bbsl) and tick-borne encephalitis virus (TBEV) in Western Europe, both pathogens are transmitted by *Ixodes ricinus* tick, therefore sharing similar vertebrate hosts [1]. Whilst small vertebrate hosts are likely to act as pathogens’ reservoir and amplifiers (such as rodents *Apodemus flavicollis* and *Myodes glareolus*) [2], larger animals like roe deer (*Capreolus capreolus*) act as tick feeding hosts for all tick stages [3,4].

In France, the presence of both pathogens is evidenced by human cases of both diseases that have been reported for several decades [5,6]. Whilst Lyme cases in humans are reported in almost all regions of France, hotspot areas are the northeastern, eastern and central parts of the country [7]. Regarding TBE, human cases have been reported every year in northeastern France, and has spread further south as evidenced by the recent cases in Alpine region and central part of the country [8–10]. Here, we hypothesize that the spatial distribution of human cases for both diseases results from several underlying spatial processes, spanning from the presence of the pathogen in the wild (in ticks or in wild vertebrate hosts), to the presence of humans and contacts with infected ticks, as well as further anthropogenic factors related to the natural history of diseases, medical consultation, and disease reporting.

Serology is a useful way to investigate the prevalence of pathogen exposure at a large scale [11] and roe deer, as sentinel hosts - defined as hosts reflecting pathogen’s dynamics with easy access for sampling [12] -appear to be ideal for tracking the presence and distribution of Bbsl and TBEV in the wild for several reasons. First, roe deer, highly infested by ticks, are common feeding hosts for *Ixodes ricinus* [13]. Second, they have been reported to develop an antibody response against both pathogens, even if the species is not a competent host for Borrelia [14–18], and has been considered as sentinel hosts for disease surveillance elsewhere [19–21]. Third, although they are widely distributed in the wild, their spatial behavior is restricted to a relatively stable area (home range) [22] allowing their infection status to represent a local spatial process. Finally, sampling is facilitated in some settings through hunting activities.

Here, to get a better insight on the distribution of Bbsl and TBEV presence in the wild, we explored the spatial pattern of roe deer serological status against both pathogens over large areas in Northern France, where *Ixodes ricinus* and at least Bbsl is known to be present as evidenced by Lyme serosurveys in forestry workers [9], and where in some areas of Northeast France TBE has been reported in humans [8], rodents or ticks [23,24].

## Material and methods

### Data collection

A total of 3,145 dried blood samples (DBS) from roe deer were collected on Whatman^®^ filter paper through hunters in 18 French departments (Figure 1AC), namely Ain, Aube, Calvados, Jura, Marne, Meurthe-et-Moselle, Meuse, Oise, Orne, Bas-Rhin, Haut-Rhin, Savoie, Haute-Savoie, Seine-et-Marne, Somme, Vosges, Essonne and Val-d’Oise, covering areas at higher risk of Borrelia and tick-borne encephalitis virus infections in humans [5,9,25]. Since information on roe deer population size is unavailable, our sampling design aimed at testing animals throughout the study area, with a minimum of 3 animals per 64 km^2^. Data were collected during three hunting seasons (from September to February), from 2019 to 2022. Information such as age, sex, location of animals, and number of observed ticks (adults and nymphs) on ear were recorded [26].

**Figure 1.**
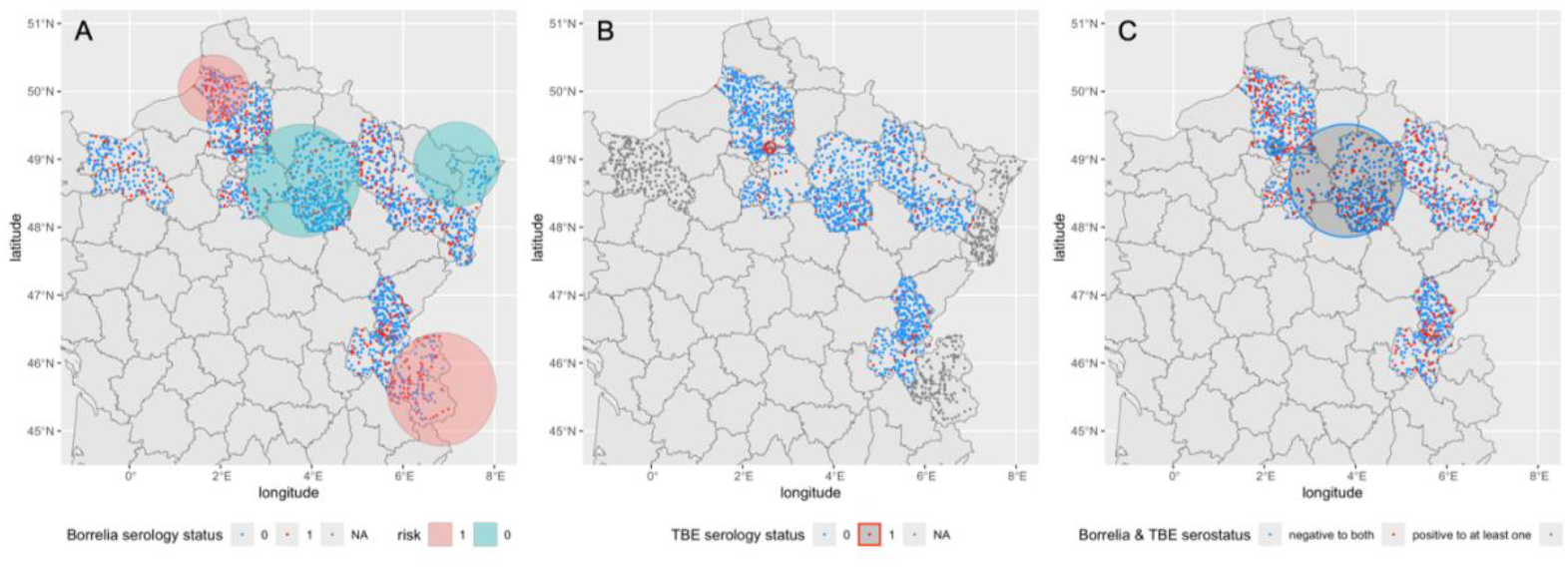
**Location of sampled animals with their serological status (red-positive, blue-negative), and results of the cluster detection tests;** for (A) Borrelia seropositive deer, (B) TBEV seropositive deer, and (C) deer seronegative to both ELISA Borrelia and TBEV tests. Grey dots indicate locations for which no ELISA tests were performed. Details of detected clusters are available in Table 3.

## Detection of anti-Borrelia and anti-TBEV antibodies

To detect antibodies against *Borrelia burgdorferi sensu lato* (Bbsl), we used a commercial anti-Bbsl ELISA kit (Borrelia + VlsE IgG ELISA Kit; IBL International) designed for human serum, that we modified and adapted to quantify the level of antibodies against Bbsl in deer serum. Details are available from Ollivier et al. 2023 [14]. Briefly, eluates were obtained by putting 1 cm^2^ of the DBS during twelve hours in a solution on a stirrer. To confirm the Borrelia serological status, we performed immunoblots (Anti-Borrelia plus VlsE Western blot IgG, EuroImmun) on 28 samples with optical density (OD) values spread over a range of values representative of the entire dataset. Western blot was used to detect specific markers present on the surface of Bbsl (specifically VlsE, OspA and OspC) to confirm the specificity of the ELISA results. As the immunoblot assay was also designed for human serum, we similarly replaced the peroxidase-conjugated antibodies to human IgG with a peroxidase-conjugated antibody to white-tailed deer IgG (Peroxidase labeled Antibodies to deer IgG (H+L); 04-31-06, KPL).

To detect anti-TBEV antibodies, we used competition ELISA Immunozym FSME IgG all species (Progen, GmbH, Germany) on DBS samples eluted in phosphate buffer saline solution (see SI Methods for details). We tested only a sub-sample of those tested against Bbsl, conditional on sufficient remaining biological material. As a consequence, samples from six departments (Bas-Rhin, Haut-Rhin, Calvados, Orne, Savoie, and Haute-Savoie) were not tested. To assess the performance of the ELISA test, 98 samples randomly chosen across the range of ELISA OD values were sent to the National Reference Laboratory for animal flaviviruses in ANSES Maisons-Alfort (France) for Virus neutralisation tests (VNT). Virus neutralization tests were conducted using the TBEV strain Hypr (Gene bank accession number U39292.1) and the microneutralization format. A sample was considered positive if the titer of neutralizing antibodies against TBEV reached the dilution 1:20.

### Threshold positivity estimations

Since we adapted ELISA kits for our study purposes, we re-estimated threshold positivity values. To do so, we first fitted a bimodal mixture model to the optical density (OD) values for both ELISA separately, and compared their fit using the Akaike Information Criteria (AIC) to a model with a single normal distribution as described in [27][27]. When the distribution of OD values was bimodal, low OD values were expected to correspond to negative samples and high OD values to positive, and the seropositivity threshold was determined with 99% confidence by the mean OD value of negative individuals plus two standard deviations, which limits false positives.

### Statistical analysis

Using the estimated threshold values, we first explored the pattern of seropositivity for both pathogens separately by estimating the overall, age-stratified and seroprevalence per department. Then, we fitted a logistic regression model to explore the factors associated with Bbsl seropositivity (sex, age, department, tick count, and TBEV ELISA status). Univariable and multivariable analyses were conducted and the best model was selected based on the AIC. Patterns of exposure to both, either or none of the pathogens at the individual level, were investigated using a multinomial logistic regression model for multiple outcomes. We assume that the different outcomes reflected a spectrum in exposure levels to infectious tick bites, in which being seronegative to both pathogens lies at the lowest bound, being seropositive to both in the upper bound, and being positive to only one pathogen lies in the middle. Finally, we searched spatial clusters of higher-risk and lower-risk of LB and TBE seropositive animals, or both. Cluster detection tests were performed using a spatial permutation Bernoulli model using Kulldorff’ scan statistic [28], implemented in SatScan 10.2 software [29]. The spatial Bernoulli model identifies areas where seropositive animals occur significantly more (high-risk clusters) or less (low-risk clusters) frequently than what is expected if those positive were uniformly distributed across locations. A circular spatial window was used, with the maximum possible size set at 50% of the positive.

### Ethics and approvals

Samples were collected as part of hunting activities following a protocol sent to local hunting groups. Data were anonymized. No specific ethical approval was required. The authors assert that all procedures contributing to this work comply with the ethical standards of the relevant national and institutional guides on the care and use of laboratory animals.

## Results

### Sampled animals

Anti-Bbsl antibodies detection test on DBS was performed on 3,145 roe deer from 18 departments (3,100 with GPS coordinates). From those 3,145 tested animals, 21.91% (N=689) were less than one-year old, 54.47% (N=1713) were estimated above 1-year old, whilst age information was not recorded for 743 animals. Females and males represented 35.26% (N = 1109) and 42.32% (N = 1331) of the Bbsl tested animals, respectively, whilst sex information was missing for 22.42% (N = 705) of the sampled animals. Anti-TBEV antibodies ELISA were performed on a subset of 2,227 samples from 12 departments (2,199 with GPS coordinates). A quarter of samples were from animals below 1-year old (25.15%, N = 560), and 63.58% (N = 1416) from animals above 1-year old (age information was missing for 254 animals, 11.41%). Females represented 39.78% (N = 886) of the animals tested, and 48.94% (N = 1090) were males (no sex information recorded for 11.27% of the tested animals, N = 251). Tick count per ear ranged from 0 to 102, and were skewed with a median at 3 (interquartile range, IQR [1-7]) (Figure S1).

### Optical densities (OD) and estimated positivity threshold values

The distributions of the OD values of both ELISA tests conducted are presented in Figure 2. For both pathogens, the mixture models which included two distributions had lower AIC than the normal models, allowing to discriminate between possible seropositive and seronegative roe deer. The resulting estimated threshold values with 99% confidence were estimated at 1.93 and 0.25, respectively for Bbsl and TBEV (Table 1). Viral neutralisation tests (VNT) to search for anti-TBEV neutralising antibodies was performed on 88 samples (out of the 98 sent for analysis) due to sample quality reasons, and resulted in 15 positive, 66 negative, and 7 non interpretable results, corresponding to a sensitivity and specificity at 60% and 61%, respectively (Figure S2, Table S1). At least one positive sample by VNT was found in 8 out of the 12 departments tested (blue stars, Figure 3), and they were spread out across the study area.

**Table 1.** AIC for normal and mixture models, and estimated threshold OD values for Borrelia and TBE serological tests.

|  | **Borrelia** | **TBE** |
| --- | --- | --- |
| **Akaike Information Criterion** |  |  |
| Normal model | 8601.103 | -930.6814 |
| Mixture model | 7849.313 | -5146.516 |
| **Threshold OD values** |  |  |
| 95% confidence | 1.533255 | 0.1977138 |
| 99% confidence | 1.932513 | 0.2454391 |

**Figure 2.**
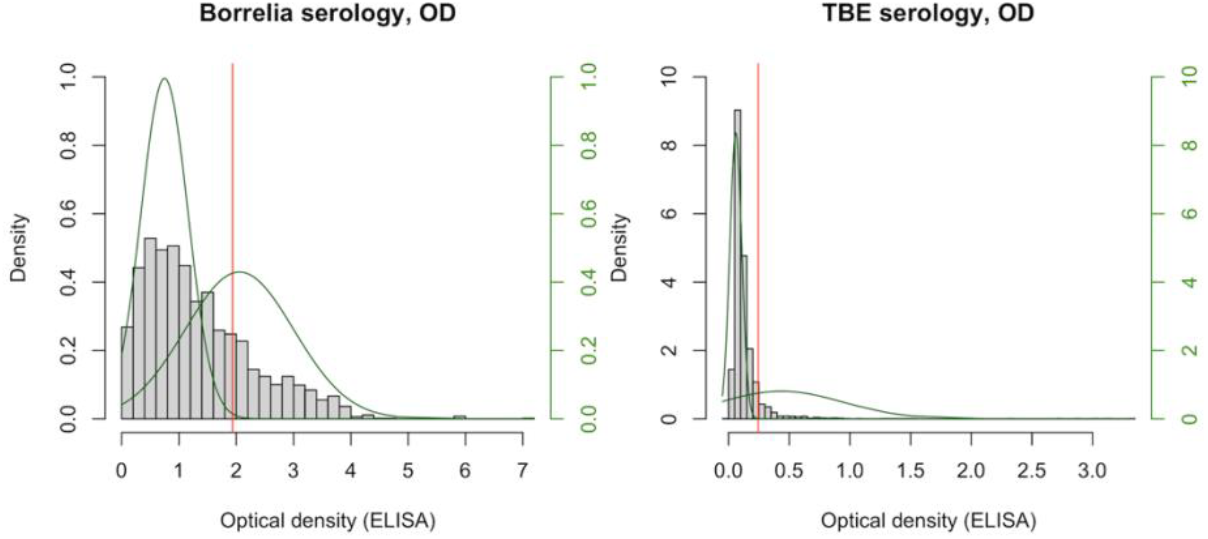
Positivity threshold estimation. Histogram of the optical density (OD) values (grey bars) and 99% confidence positivity thresholds (vertical red lines) for (A) Bbsl and (B) TBEV antibody detection. The green lines show the probability density functions of the two normal distributions fitted to the bimodal mixture model.

**Figure 3.**
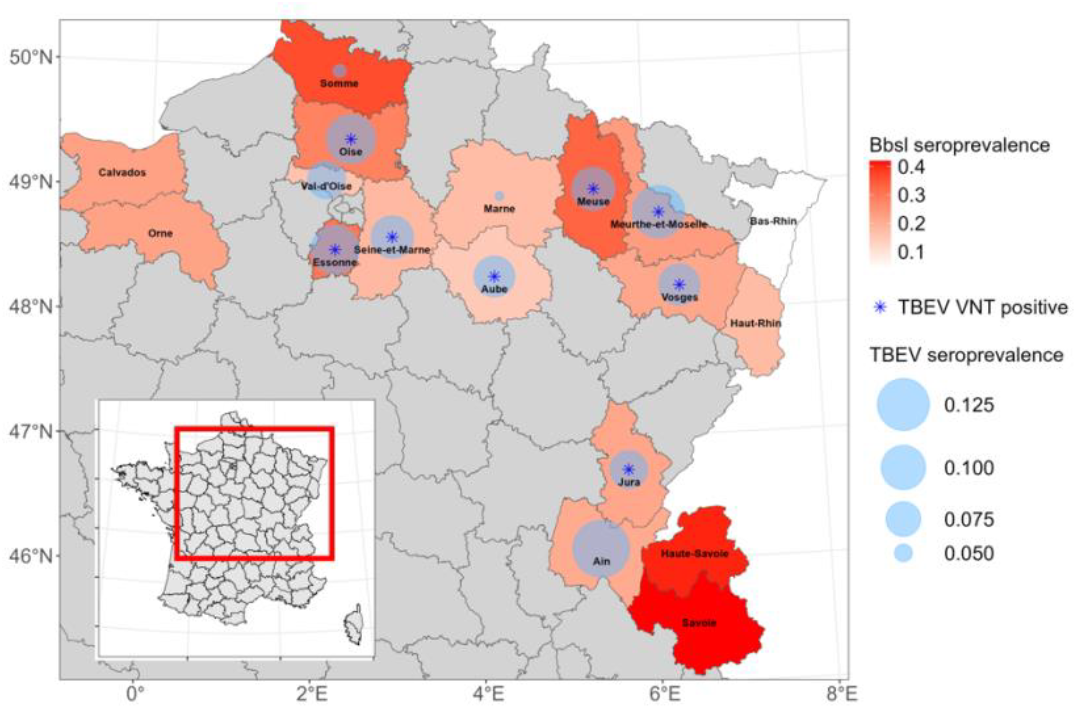
**Map showing the mean ELISA seroprevalence per department** against Bbsl (red shades) and TBEV (blue circle size). Blue stars pinpoint departments in which at least one animal was found positive to TBEV VNT. 95%CI are shown in Figure 4.

### Seroprevalence estimates and logistic regression

Using the 99% confidence threshold values, the overall ELISA IgG prevalence was 2.7 times higher for Bbsl than for TBEV, with 23.22% for Borrelia (95%CI [21.75-24.70], n=730, N=3,145) and 8.51% for TBE (95%CI [7.36 - 9.67], n=189, N=2,227). Seroprevalence in animals under or over one-year old were similar for both pathogens (Figure S3). Figure 3, Figure 4 and Table S2 present ELISA IgG prevalence results against Bbsl and TBEV, by department. The highest anti-Bbsl seroprevalence were reported in Savoie (41.92%, 95%CI [32.32-51.61]) and Haute-Savoie (40.44%, 95%CI [28.98-52.05]) both in the Alpine area, followed by Somme (36.23%, 95%CI [30.29-42.24]) and Meuse (32.89%, 95% CI [27.27-38.59]), being in the Northern and Eastern parts of the country. Most of the other departments had Bbsl seroprevalence average values ranging between 15% and 30%, whilst Bas-Rhin reported only 4.02% of seropositive animals (95%CI [0.06-8.15]). TBEV seroprevalence reached its highest value in Ain at 14.34% (95%CI [8.81-20.14]), followed by Meurthe-et-Moselle at 12.64% (95%CI [6.22-19.53]), Oise and Essonne at about 11% yet with overlapping confidence intervals. The lowest TBEV seroprevalences were reported in the Marne and Somme with respectively 4.5% (12/267) and 5.0% (11/242). At the individual level, all departments accounted for, being Bbsl seropositive was positively associated with being TBEV seropositive, with animals seropositive to Bbsl having almost twice the odds of being seropositive to TBEV than seronegative Bbsl animals (OR =1.84, 95%CI [1.34-2.53]) (Table 2). The results of the logistic model showed furthermore that this association was significant in five departments, and highest in those reporting highest Borrelia seroprevalence (mean odds ratio OR_Somme_=3.46, OR_Meuse_=2.81, OR_Essonne_=2.34, OR_Oise_=2.27, OR_Jura_=1.59), and that animals with more than one observed tick attached to the ear were at increased risk of being seropositive to Bbsl (Table 2). Despite the absence of significant interaction between TBEV sero-status and department on the Bbsl serological status in the logistic model, we investigated further the co-exposure status in those departments, by fitting a multinomial logistic regression, with four outcomes levels. Only in three departments (Somme, Meuse, Oise), the odds of animals being positive to both pathogens or only Borrelia positive, was higher than being seronegative to both pathogens, whilst the presence of more than one attached tick seen on the ear was associated with being positive to either pathogen or both (Figure S4 and Table S4),

**Table 2.** **Results of the logistic model** (odds ratio, 95% CI, *p*-value) on Bbsl serological status

| Variables | OR | 95% CI | *p*-value |
| --- | --- | --- | --- |
| **ELISA TBE serostatus** |  |  |  |
| Negative | ref | - | - |
| Positive | 1.84 | 1.32-2.55 | < 0.001*** |
| **Departments** |  |  |  |
| 10-Aube | ref | - | - |
| 80-Somme | 3.46 | 2.26-5.38 | < 0.001*** |
| 55-Meuse | 2.81 | 1.83-4.35 | < 0.001*** |
| 91-Essonne | 2.34 | 1.12-4.71 | 0.019* |
| 60-Oise | 2.27 | 1.44-3.61 | < 0.001*** |
| 39-Jura | 1.59 | 1.03-2.47 | 0.039* |
| 54-Meurthe-et-Moselle | 1.54 | 0.83-2.82 | 0.163 |
| 01- Ain | 1.44 | 0.83-2.45 | 0.188 |
| 88-Vosges | 1.42 | 0.86-2.36 | 0.168 |
| 51-Marne | 1.32 | 0.83-2.11 | 0.246 |
| 77-Seine-et-Marne | 1.22 | 0.69-2.11 | 0.482 |
| 95-Val d'Oise | 1.04 | 0.46-2.15 | 0.922 |
| No. ticks observed on ear |  |  |  |
| <= 1 | ref | - | - |
| >1 | 1.33 | 1.07-1.66 | 0.010 |

**Table 3.**
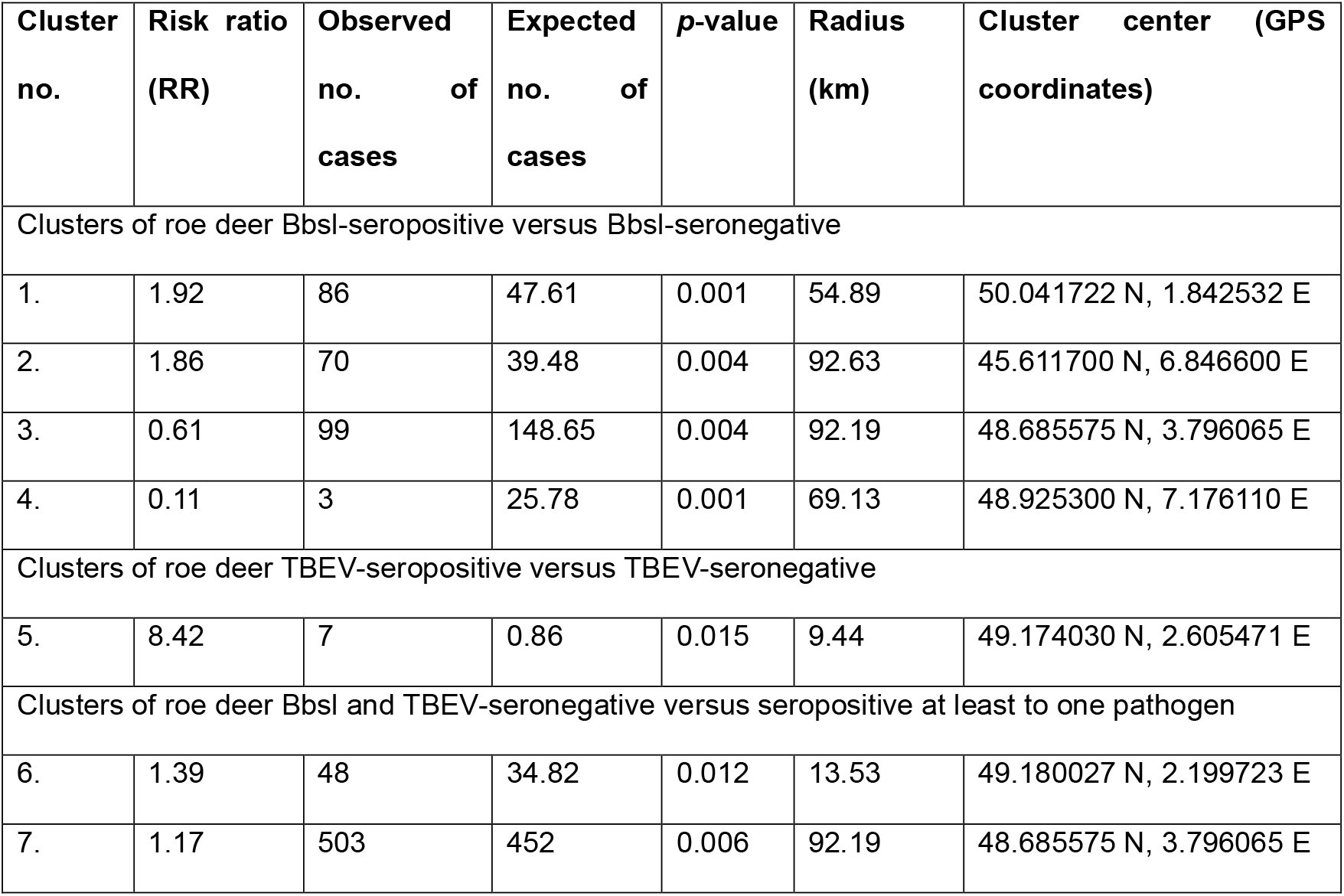
**Results of the cluster detection tests**, of Bbsl seropositive, TBEV seropositive, and Bbsl and TBEV seronegative animals

**Figure 4.**
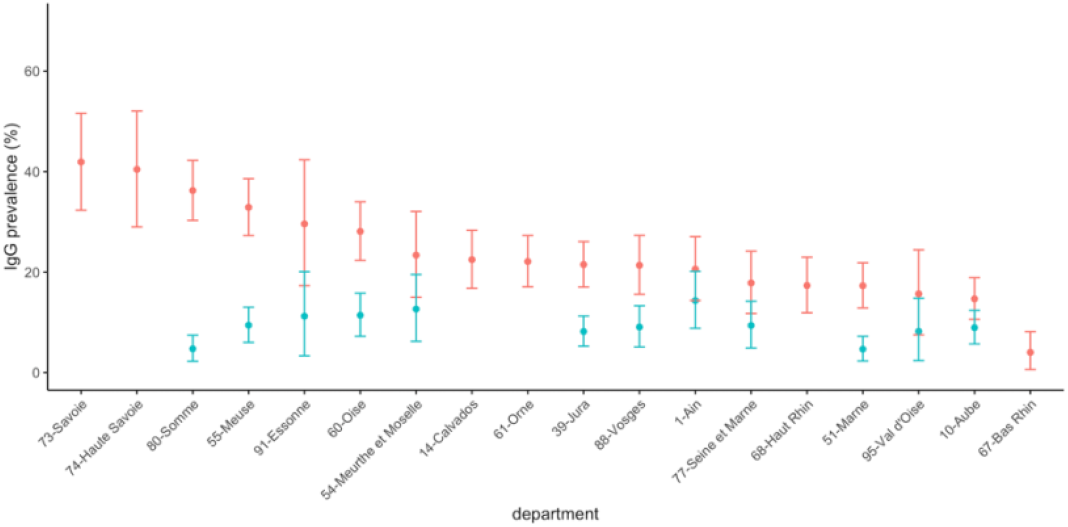
**Mean ELISA seroprevalence** against Bbsl (red dots) and TBEV (blue dots) per department, and 95% CI

### Cluster detection tests

We identified a total of seven spatially significant clusters (Table 3). Two were at high-risk and two were at low-risk of Bbsl seropositive animals. One high-risk cluster was located in the Alps region, covering some large parts of Savoie and Haute-Savoie (cluster 2, RR=1.86, radius=92.63 km, *p*-value=0.004); whilst a second cluster of smaller size was centered in the Somme department (cluster 1, RR=1.92, radius=54.89 km, *p*-value=0.001) in the northern part of the country (red circles, Figure 1A). Bbsl low-risk clusters were located in the center (cluster 3, RR=0.61, radius=92.19km *p*-value=0.004) and northeastern part of the country (cluster 4, RR= 0.11, radius 69.13km, *p*-value=0.001), respectively (blue circles, Figure 1A). In contrast, the distribution of TBE positive animals did not exhibit any strong pattern, with only one small size high-risk cluster detected in the Oise department (cluster 5, RR = 8.42, radius=9.44km, *p*-value=0.015) (Figure 1B). Finally, individuals with both negative statuses were significantly clustered in two areas, with the largest cluster (cluster 7) (Figure 1C), overlapping with the largest low-risk Borrelia cluster (cluster 3).

## Discussion

We present a serological study conducted in roe deer over a large sampled area in Northern France, where *Ixodes ricinus* is known to be present, and covering areas where Lyme borreliosis and/or TBE cases have been reported in humans. ELISA seropositive roe deer against both pathogens were identified across the investigated departments, and VNT TBEV positive animals were found in most of these departments. Two spatial clusters at higher Borrelia seropositivity were detected, whilst no marked spatial pattern reported for TBEV exposure.

Similar levels of Bbsl and TBEV seroprevalence were reported in the few studies conducted in the neighbouring areas, over the past 25 years. Two ELISA studies conducted in roe deer on both pathogens were reported, from Denmark with 36.6% and 8.7% seropositive for Bbsl and TBEV [30], respectively, and more recently in northern Italy with 8.8% and 26.6%, for Bbsl and TBEV seroprevalence [31]. Nevertheless, TBEV seroprevalence was more investigated than Bbsl in our study hosts. Reported TBEV IgG prevalence was assessed with various techniques, ranging from 2% in The Netherlands in 2010 using VNT [32] to 22.9% (24/105) or 22.86% (8/35) in two studies in Germany between 2007 and 2013 using the similar ELISA Immunozym FSME IgG we used [33], or VNT [20] [20]. In-between estimates were reported in Austria with 2.32% seropositivity (among 945 tested using Immunofluorescence Antibody Test (IFAT) [19], 3.4% in The Netherlands (among 640 animals tested using Immunozym FSME IgG, with VNT on positive and bordeline samples) over a seven years period [34], 4.07% in the UK in 2018-2019 using also the same Immunozym FSME IgG All Species kit [35], 5.10% in Flanders (98 animals tested by VNT, 2008-2013) [36], 6.9% in Denmark in 2013-2014 (51/736 based on VNT) [37], and 12.4% in Belgium in 2007-2009 using Immunozym FSME IgG [38].

In France, spatial information on Bbsl and TBEV from animal populations are limited. The sole study exploring Bbsl seroprevalence in roe deer was conducted on 765 samples collected between 1979 and 1988. From the 12 departments tested across the country, only two departments (Haute-Marne and Meurthe-et-Moselle) had overall 35.5% of seropositive animals (35.5%, 27/76) [39], the latter department reporting over 20% seropositive animals in our study. More recently, very low seropositivity against anti-TBEV-like antibodies was found in roe deer in southwestern France (0.13%), using a Flavivirus ELISA kit followed by microsphere immunoassays and micro-neutralization tests [40]. Regarding human data, Lyme disease and TBE cases reported through national surveillance, and seroprevalence surveys in forestry workers, reported higher-risk areas in the northeastern part of the country, in the Alpine area, and in the Central region of the country to a lower extent [5,6,9]. Our data similarly exhibited a Bbsl high-risk cluster in Alpine region (Savoie, Haute-Savoie), and a high-risk cluster in northern France (Somme department) where Lyme disease is regularly reported in humans. However, whilst we would have expected to detect a high-risk Bbsl seropositive cluster in Alsace-Lorraine, our data exhibits the opposite, that is, a low-risk significant cluster. Whilst we do not have any epidemiological explanation for these results, we assumed a bias may have occurred in that area during animal sampling or sample testing. Regarding TBEV, no marked pattern was evidenced, yet the highest seropositivity was identified in Ain department, where the cluster of 44 food-borne human cases was reported in 2020 [10]. Our data exhibited a small high-risk TBEV seropositivity cluster in the Oise department (northern Paris), from where no other evidence of TBEV presence had been reported.

The absence of detection of smaller LB clusters or TBEV exposure clusters may be related to unsufficient sampling efforts at lower resolution. However, our uniform sampling design over a grid allowed to investigate the presence of pathogens over large areas. Future studies should target sampling more animals in those specific high-risk and low-risk areas, and expand spatial sampling over the whole country, including areas at lower risk (southern part of the country) to provide key baseline information to interpret further those results.

Seroprevalence studies are powerful designs to assess past or recent exposure to pathogens [11]. Yet, their accurate interpretation relies on knowledge on antibodies persistence following exposure to pathogens, and the performance of ELISA tests (sensitivity and specificity) in target animal species. There is limited knowledge on the persistence of antibodies against Bbsl and TBEV in animals. Whilst anti-TBEV antibodies may persist lifelong in humans [41], they could be detected several months following infection in wild voles in controlled laboratory settings [42]. For Bbsl, antibody persistence is yet to be clarified in humans [43], and has been estimated to only a few weeks using capture-recapture data in roe deer [14]. Serological analyses looking at anti-TBEV antibodies in samples from long-term capture-mark-recapture monitoring data in roe deer populations, such as those used to explore the temporal persistence of anti-Bbsl antibody levels in two French populations [14], would provide valuable information in this context. Moreover, ELISA tests commercially available are not developed to detect Bbsl nor TBEV antibodies in our studied species. However, we maximized the use of those commercially available tests by using secondary antibodies in the Borrelia kit, by conducting VNT to assess the TBE kit, and by re-estimating threshold values. Furthermore, whilst the sensitivity and the specificity of TBEV ELISA kit were moderate, they aligned with recent estimates yielded from cattle [23], and they are the first estimates targeting roe deer. Indeed, previous studies using similar kits confirmed the status only of positive or doubtful samples [33–35,38]. Corrected ELISA seroprevalence from our estimated sensitivity and specificity would yield very low IgG TBE seroprevalence, yet VNT results suggest the presence of anti-TBEV antibodies across the study area. Our study offers valuable insight into the need to develop better ELISA test for the pathogens and targeted species.

This large serosurvey in roe deer in Northern France suggests evidence of TBEV and Bbsl antibody presence across a large study area. Further research would benefit from increasing sampling efforts locally, testing from areas located further South to get a better insight on the large-scale spatial pattern, and improve serological tests for the use in roe deer.

## Acknowledgements

Authors wish to thank all participating hunters and departmental laboratories for the sampling and data collection.

## Authors contributions

Conceptualization: RM, TB

Data curation: RM, MB, VO, AP, BC

Formal analysis: RM, VO, AP, IES

Funding acquisition: RM, MB, BC, TB

Investigation: RM, MB, VO, AP, BC, GG, CM, TH, IES, YK

Methodology: RM, MB, TB

Project administration: RM, MB, BC, HV, TB

Resources: RM, TB, MB, BC, GG

Software

Supervision: RM, MB, BC, HV, TB

Validation

Visualization: RM, IES

Writing – original draft: RM

Writing – review & editing : RM, MB, VO, AP, BC, GG, TH, CM, IES, YK, HV, TB

